# Cardioluminescence in transgenic zebrafish embryos: a Ca^2+^ imaging tool to study drug effects and pathological modeling

**DOI:** 10.1101/2021.05.06.442917

**Authors:** Manuel Vicente, Jussep Salgado-Almario, Michelle M. Collins, Antonio Martínez-Sielva, Masafumi Minoshima, Kazuya Kikuchi, Beatriz Domingo, Juan Llopis

## Abstract

The zebrafish embryo has emerged as an excellent model in cardiovascular research. The existing techniques to monitor Ca^2+^ in the heart based on fluorescent Ca^2+^ biosensors are limited due to phototoxicity and photobleaching. To overcome these issues, we have used bioluminescence. We generated a transgenic line expressing GFP-Aequorin in the heart, *Tg*(*cmlc2:GA*), and optimized an *in vivo* aequorin reconstitution protocol to improve the luminescence capacity. This allowed imaging Ca^2+^ in long duration recordings in embryos of 3 to 5 days post-fertilization. The analogs *diacetyl h*-coelenterazine and *f*-coelenterazine enhanced the light output and signal-to-noise ratio from the embryos. With this cardioluminescence model, we monitored the time-averaged Ca^2+^ levels and beat-to-beat Ca^2+^ oscillations. Changes in Ca^2+^ levels were observed by incubation with BayK8644, an L-type Ca^2+^ channel agonist, the channel blocker nifedipine, and β-adrenergic blocker propranolol. Treatment of zebrafish embryos with terfenadine for 24 hours has been proposed as a model of heart failure. *Tg*(*cmlc2:GA*) embryos treated with terfenadine showed a 2:1 atrioventricular block and a decrease in the ventricular Ca^2+^ levels.

## 1. Introduction

Cardiovascular diseases are the leading cause of death worldwide and are among the most challenging to diagnose and treat due to the complexity of their pathophysiology. Ca^2+^ plays a pivotal role in the excitation-contraction coupling of the heart and alterations in Ca^2+^ cycling, its associated proteins and pathways may trigger pathological disorders [1, 2]. Ca^2+^ handling has been extensively studied, mainly in isolated cardiomyocytes obtained from animals or from differentiated human pluripotent stem cells [3–6]. While this approach provides a detailed insight of the Ca^2+^ fluxes, the interaction between the heart, other organs and the nervous and endocrine systems is lost. Thus, to better understand the underlying pathological processes, *in vivo* experimental (animal) models are crucial. The zebrafish embryo has emerged as an excellent model in cardiovascular research because of its high genetic manipulability, ex-utero embryonic development and transparency [7–13]. The zebrafish heart is composed of only two chambers, an atrium, and a ventricle. Despite this anatomical difference, many studies have argued that zebrafish heart physiology, heart rate (HR) and action potential resemble those of the human [14–16], although some of the ionic channels and regulation are not identical [17–18].

Non-invasive Ca^2+^ imaging techniques in zebrafish embryos may contribute to our understanding of Ca^2+^ handling and its relationship with cardiac diseases. Genetically encoded Ca^2+^ biosensors, like the single-fluorophore GCaMPs, have been successfully employed to image Ca^2+^ dynamics in the embryonic heart [11, 19–21]. This requires disrupting cardiac mechanical function with morpholino oligomers against myosin II [22] or with inhibitors like *para*-amino blebbistatin [23] to avoid motion imaging artifacts. We have previously used ratiometric biosensors, which require special optical components to acquire two emission images simultaneously [24], to correct these artifacts. Although fluorescent biosensors provide good temporal resolution of the Ca^2+^ transients, the continuous illumination may cause phototoxicity and photobleaching of the biosensor, limiting the duration of an imaging experiment to a few seconds, depending on the instrumentation. A solution is the use of low-light-level techniques like light sheet microscopy, much less harmful to the cells [20, 25]. However, a further pitfall is the autofluorescence of the vitello, adjacent to the heart. By contrast, bioluminescence imaging does not need excitation light and has been robustly demonstrated to be biocompatible in this animal model [26–29]. The photoprotein aequorin has been widely employed to measure Ca^2+^ in many cell types and animal models [30–32]. The functional photoprotein is formed when apoaequorin binds the substrate coelenterazine (CTZ) [30]. In our previous work, aequorin fused with GFP (GA) [33] was used to visualize cytoplasmic and mitochondrial Ca^2+^ in skeletal muscle of zebrafish embryos for hours [29]. Despite its low photon yield, we hypothesized that GA could be used to measure the synchronized Ca^2+^ transients in the heart. However, it might be challenging to achieve sufficient reconstitution with CTZ in the presence of constant Ca^2+^ cycling, since aequorin is consumed once it emits light.

In this study, we have generated a transgenic zebrafish line expressing GA, *Tg*(*cmlc2:GA*), to image continuously ventricular Ca^2+^ levels *in vivo* (cardioluminescence). Ca^2+^ was measured as the heart was performing its mechanical function in 3-, 4- and 5-days post-fertilization (dpf) zebrafish embryos. To improve the efficiency of the reconstitution of aequorin with CTZ, embryos were treated with the L-type Ca^2+^ channel (LTCC) blocker nifedipine to decrease temporarily Ca^2+^ cycling.

## 2. Results

### 2.1 Tg(cmlc2:GA) zebrafish line generation

We generated a transgenic zebrafish line expressing GA under the control of the cardiac-specific promoter *cmlc2*. Stable expression of GA did not affect embryo survival. In this chimera, upon Ca^2+^ binding to aequorin there is energy transfer between the excited state of CTZ and the GFP, so that light emission is shifted to the green (Fig. 1A) [31, 33, 34]. The expression of GA, reported by the fluorescence of GFP, was observed at 24 hours post-fertilization when the cardiac tube was formed, and no fluorescence was seen in other organs (Fig 1C). The ventricle was brighter than the atrium at 3, 4 and 5 dpf, likely due to the thickness of its wall (Fig. 1B). Both the HR and the ventricular fractional shortening of heterozygous GA embryos were like those observed in wild-type siblings (Fig. 1D and E).

**Fig. 1.**
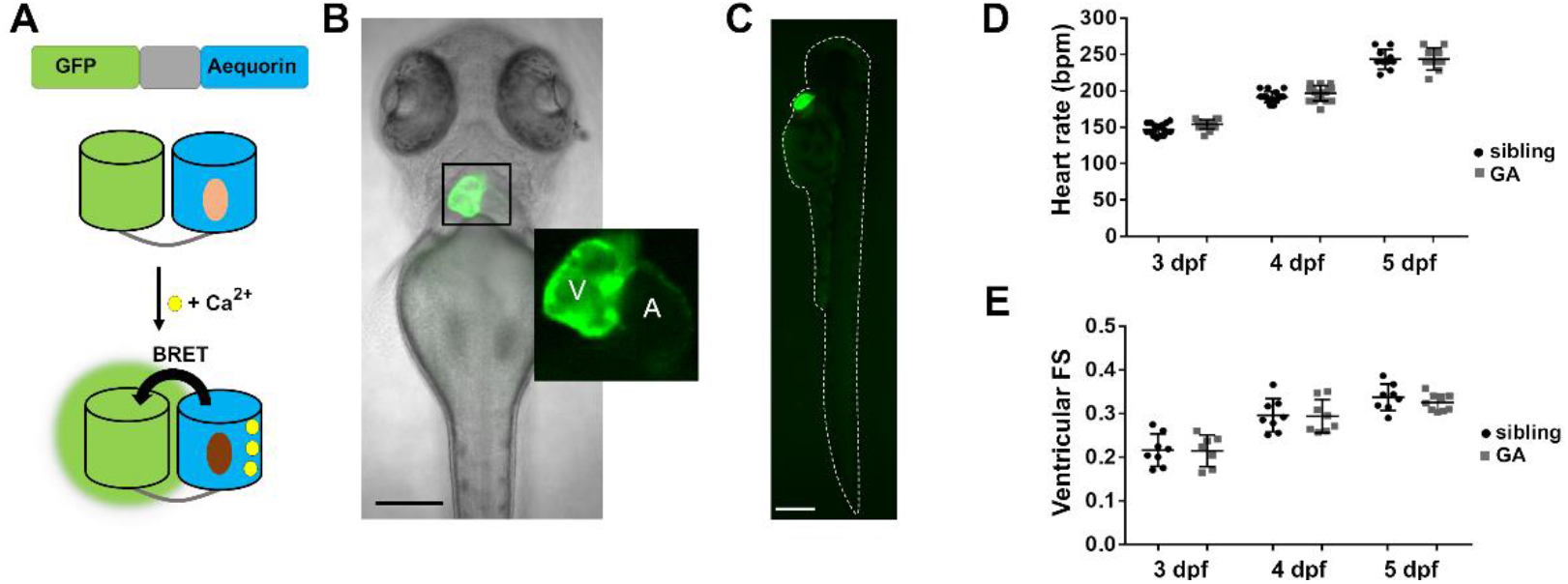
Expression of GA in the heart of zebrafish embryos. **A)** DNA construction and concept of the Ca^2+^-dependent bioluminescence of the chimera GA. BRET, bioluminescence resonance energy transfer. Light and dark brown ovals represent CTZ and its excited product coelenteramide, respectively. **B)** Superimposed GFP fluorescence and transmitted light images of a 3 dpf *Tg*(*cmlc2:GA*) zebrafish embryo. The inset shows the atrium (A) and ventricle (V). **C)** GFP fluorescence image of a 3 dpf *Tg*(*cmlc2:GA*) zebrafish embryo. **D)** HR of GA heterozygous and wild-type sibling embryos at 3, 4 and 5 dpf. A two-tailed unpaired *t*-test was used. Data are shown as the mean ± SD (sibling n=17 for 3 and 4 dpf and n=10 for 5 dpf; GA n=17 for 3 and 4 dpf and n=10 for 5 dpf). **E)** Ventricular fractional shortening (FS) of GA heterozygous and sibling embryos at 3, 4 and 5 dpf. A two-tailed unpaired *t*-test was used. Data are shown as the mean ± SD (sibling n=8 for 3, 4 and 5 dpf; GA n=7 for 3, n=8 for 4 dpf and n=9 for 5 dpf). No statistical differences were found in D and E (p > 0.05). Bar scale indicates 150 μm and 250 μm for B and C, respectively.

### 2.2 Aequorin reconstitution protocol

The apoaequorin needs to be reconstituted with CTZ in the presence of oxygen to yield the Ca^2+^-sensitive photoprotein [35]. However, when 3 dpf *Tg*(*cmlc2:GA*) zebrafish embryos were incubated in 50 μM *diacetyl h*-CTZ for 2 hours, spontaneous Ca^2+^-dependent bioluminescence from the heart was not detected (Fig. 2A). Triton-X100 (5%), added to bring aequorin into contact with extracellular Ca^2+^ to quantify total luminescence, released few counts (Fig. 2A) suggesting that too little aequorin was reconstituted. As Ca^2+^ rises in each systole, the rate of aequorin consumption might be faster than the reconstitution rate. Therefore, we hypothesized that limiting Ca^2+^ transients during aequorin reconstitution would improve light output.

**Fig. 2.**
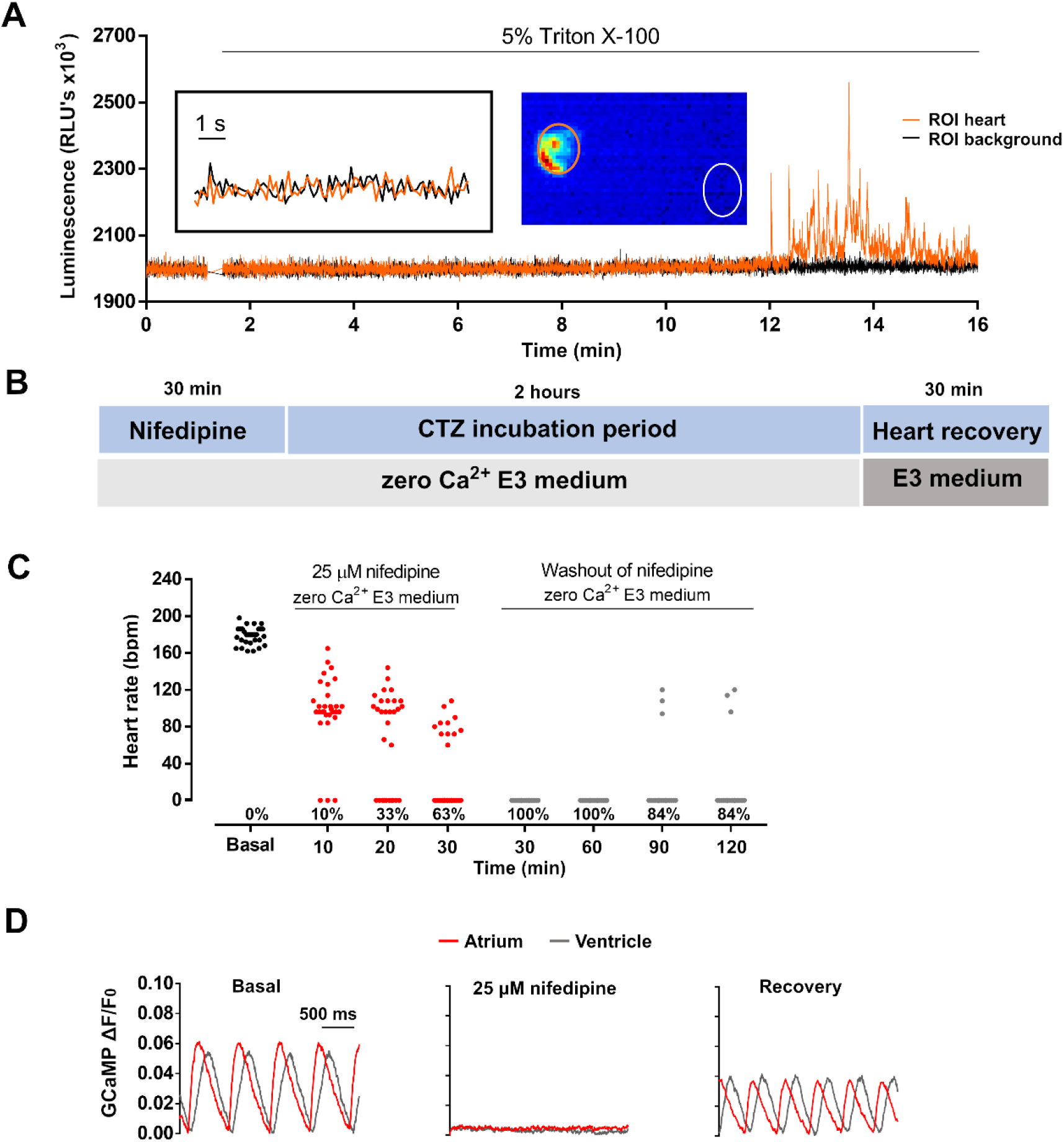
Limiting Ca^2+^ transients in the heart with nifedipine and aequorin reconstitution protocol. **A)** Representative luminescence recording acquired at 9 Hz of a beating heart in a 3 dpf *Tg*(*cmlc2:GA*) zebrafish embryo incubated with 50 μM *diacetyl h*-CTZ for 2 hours. The image represents the integrated luminescence of the entire experiment. Regions of interest (ROI) from heart and background are shown. **B)** Scheme of the aequorin reconstitution protocol comprising a 30-min incubation with nifedipine in zero Ca^2+^ E3 medium to block Ca^2+^ transients, 2 hours for aequorin reconstitution with CTZ, and 30 min for recovery of the Ca^2+^ transients in Ca^2+^-containing E3 medium. **C)** HR of 3 dpf *Tg*(*cmlc2:GA*) zebrafish embryos in basal conditions and during the treatment with 25 μM nifedipine (n=30). Only the non-beating embryos at the end of the nifedipine treatment (n=19) were tracked during the washout period in zero Ca^2+^ E3 medium (*CTZ incubation period* in B). The % of non-beating embryos at each time is indicated. **D)** Representative Ca^2+^ transients of 3 dpf *Tg*(*cmlc2:GCaMP*)^*s878*^ embryos in basal conditions, treated with 25 μM nifedipine for 30 min in zero Ca^2+^ E3 medium and after the heart recovery period. The fluorescence images were acquired at 200 Hz.

#### 2.2.1 Suppressing Ca^2+^ rise in the heart with an LTCC blocker

A protocol was devised to blunt Ca^2+^ transients by incubating embryos with 25 μM nifedipine in zero Ca^2+^ (Fig. 2B). Suppl. Fig. 1 shows the optimization of this treatment. After 30 min in nifedipine, non-beating embryos were transferred to a plate containing zero Ca^2+^ E3 medium for 2 hours. This period will be used for aequorin reconstitution with CTZ (Fig. 2B). Then, embryos were placed in complete E3 medium for 30 min to allow for heart recovery. In the *Tg*(*cmlc2:GA*) line, the heart was completely stopped by the treatment with nifedipine in 63 % of the embryos and 84 % of those remained motionless after 2 hours (Fig. 2C). The Ca^2+^ levels during this protocol were monitored with the Ca^2+^ biosensor GCaMP in the *Tg*(*cmlc2:GCaMP*)^*s878*^ zebrafish line (Fig. 2D). The Ca^2+^ transients were indeed abrogated by the treatment with nifedipine and restored after the heart recovery period. This could allow sufficient aequorin reconstitution to image Ca^2+^ in the zebrafish heart.

#### 2.2.2 Embryos treated with the aequorin reconstitution protocol recovered heart function after restoring Ca^2+^ into the medium

We tested the recovery of the cardiac function in embryos subjected to the aequorin reconstitution protocol (Fig. 2B). The HR was similar before and after the aequorin reconstitution protocol in 3 dpf (170 ± 9 bpm before *vs*. 166 ± 15 bpm after p=0.387), 4 dpf (192 ± 16 bpm before *vs*. 179 ± 29 bpm after; p=0.163) and 5 dpf embryos (191 ± 16 bpm before *vs*. 181 ± 24 bpm after; p=0.071) (Suppl. Fig. 2A). In addition, the ventricular fractional shortening was restored after the recovery period at 3 dpf (0.23 ± 0.02 for control *vs*. 0.24 ± 0.02 for Aeq protocol; p=0.832), 4 dpf (0.30 ± 0.04 for control *vs*. 0.29 ± 0.02 for Aeq protocol; p=0.315) and 5 dpf (0.31 ± 0.02 for control *vs*. 0.31 ± 0.02 for Aeq protocol; p=0.878) (Suppl. Fig. 2B).

The dynamics of Ca^2+^ transients were further studied in 3 dpf *Tg*(*cmlc2:GCaMP*)^*s878*^ embryos subjected to the aequorin reconstitution protocol. As shown in Suppl. Fig. 2C and D, the Ca^2+^ transients were similar in control embryos and embryos treated with the protocol in the atrium and ventricle. Moreover, the kinetics of the Ca^2+^ transients, such as the rise time 10-90% and the decay time 90-10% (Suppl. Fig. 2 E-G), were also similar between control embryos and embryos treated with the protocol. These results suggest that the heart regained normal Ca^2+^ transients and mechanical function during the recovery period.

#### 2.2.3 Imaging individual Ca^2+^ transients in the heart

We tested the bioluminescence obtained in embryos treated with the aequorin reconstitution protocol. Two *Tg*(*cmlc2:GA*) zebrafish embryo groups were incubated in 50 μM *diacetyl h*-CTZ for 2 hours with or without the previous treatment with 25 μM nifedipine. Cardioluminescence, the spontaneous bioluminescent flashes due to beat-to-beat Ca^2+^ oscillations during the cardiac cycle, was detected only in embryos preincubated with nifedipine (Fig. 3A). We calculated the signal-to-noise ratio (SNR) of these recordings, a measure of the sensitivity of the method (see Materials and Methods). Fig. 3B shows that embryos treated with nifedipine had higher SNR than the controls at 3, 4 and 5 dpf. Triton X-100 was added to compare the amount of reconstituted aequorin in each case. Embryos preincubated with nifedipine released 5 to 18-fold more counts than those from the control group at 3, 4 and 5 dpf (Fig. 3C). Taken together, these results confirmed that limiting the Ca^2+^ transients during the incubation with CTZ improved the efficiency of aequorin reconstitution and the luminescence signal from the heart in *Tg*(*cmlc2:GA*) zebrafish embryos, such that individual ventricular Ca^2+^ transients could be observed. However, spontaneous luminescence from the atrium was not detected in most embryos (Suppl. Fig. 3 A-C). Total counts in the atrium after Triton X-100 were significantly less than those in the ventricle but were detectable (Suppl. Fig. 3D-E), showing the need to separate spatially the origin of the luminescence. In contrast to photometry, imaging allowed us to draw ROIs over the atrium and ventricle to discriminate their contribution.

**Fig. 3.**
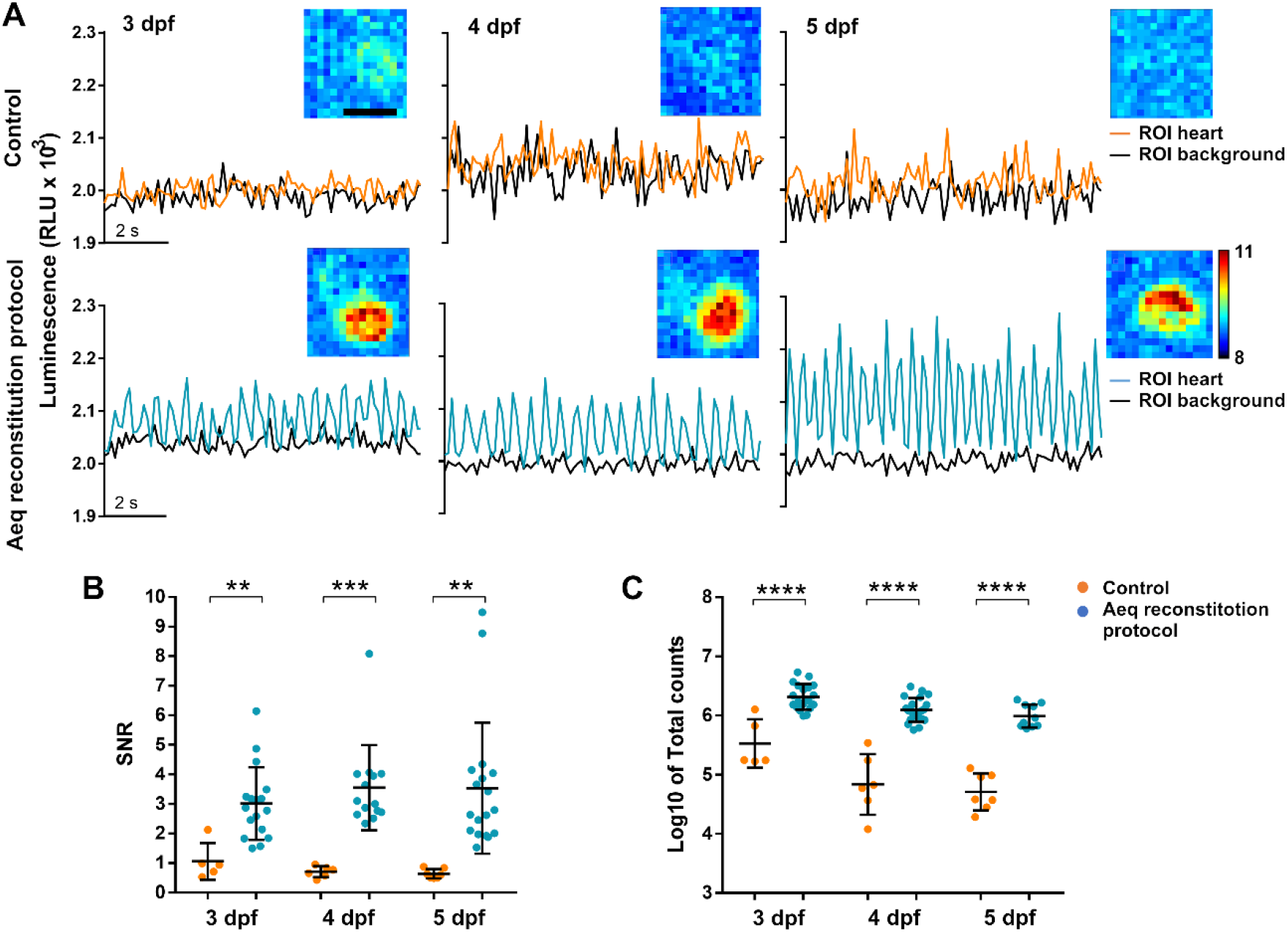
Bioluminescence imaging in the heart of 3, 4 and 5 dpf *Tg*(*cmlc2:GA*) zebrafish embryos treated with the aequorin reconstitution protocol. **A)** Luminescence recordings in CTZ-incubated embryos either with nifedipine treatment and subsequent wash-out and recovery (aequorin reconstitution protocol, Fig. 2B) or without this protocol (control). *Diacetyl h-*CTZ was used for reconstitution and images were acquired at 9 Hz. The black lines indicate the luminescence in background ROIs drawn out of the embryos. Inset images show the integrated luminescence for 1 min over the heart. The scale bar represents 50 μm and the color scale indicates RLU. **B)** SNR of the control (orange) and aequorin reconstitution protocol (blue) embryos at 3, 4 and 5 dpf. A two-tailed unpaired *t*-test was used. Data are shown as the mean ± SD (control n=5 for 3 dpf, n=6 for 4 dpf and n=7 for 5 dpf; aequorin reconstitution protocol n=17 for 3 dpf; n=14 for 4 dpf and n=18 for 5 dpf). **C)** Total counts released in the control (orange) and aequorin reconstitution protocol (blue) groups at 3, 4 and 5 dpf. A two-tailed unpaired *t*-test was used. Data are shown as the mean ± SD (control n=5 for 3 dpf, n=6 for 4 dpf and n=7 for 5 dpf; aequorin reconstitution protocol n=19 for 3 dpf; n=21 for 4 dpf and n=10 for 5 dpf). ** p < 0.01, *** p < 0.001, **** p < 0.0001.

Decreasing the acquisition frequency can improve the SNR of low emitting samples. Indeed, a strong inverse correlation was observed between SNR and the acquisition frequency (R^2^=0.911; p=0.045) (Suppl. Fig. 4A). In contrast, when luminescence signal (in relative luminescence units, RLU) is transformed into luminescence emission rate (L, in counts per second), it becomes independent of the image acquisition frequency (R^2^=0.349; p=0.409) (Suppl. Fig. 4B).

### 2.3 Testing different CTZ analogs

In our experiments, to generate functional aequorin, CTZ must diffuse through the skin and tissues to reach the heart and cross the cardiomyocyte plasma membrane. Different CTZ synthetic analogs afford varying chemical stability, Ca^2+^ sensitivity and membrane permeability. We therefore tested several CTZ analogs (Suppl. Fig. 5) to optimize the cardioluminescence from the embryos.

The Ca^2+^ sensitivity of *h*-, *f*-, *fcp*-, and *hcp*-CTZ has been shown to be 16, 20, 135 and 190-fold higher than that of *native-CTZ*, respectively [36]. The rate of reconstitution is also affected by the analog used: *native-CTZ* was 7 to 10-fold faster than *f*- and *h*-CTZ, respectively [37]. Furthermore, *f*-CTZ was reported to have nearly 2-fold more membrane permeability than *native*- and *h*-CTZ [38]. The addition of a diacetyl group to *h*-CTZ improves its chemical stability and has been shown to allow long-term imaging by providing a constant supply of substrate [39]. Therefore, we undertook its chemical synthesis (see Materials and Methods). The SNR and the total counts obtained in *Tg*(*cmlc2:GA*) embryos from 3 to 5 dpf reconstituted with *native*-, *h*-, *fcp*-, *hcp*-, *f*- and *diacetyl h*-CTZ were evaluated (Fig. 4A and B). Both *f*- and *diacetyl h*-CTZ provided the best balance for cardioluminescence experiments, with robust light output at physiological Ca^2+^ levels and high SNR. We used *diacetyl h*-CTZ in further experiments.

### 2.4 GA reports the effect of drugs acting on LTCC in the zebrafish ventricle

To validate the functionality and sensitivity of the *Tg*(*cmlc2:GA*) line, we used two drugs to target the LTCC: the agonist Bay K8644 and the antagonist nifedipine. Fig. 5A shows a cardioluminescence recording acquired at 2 Hz in a 3 dpf embryo. This lower acquisition frequency was used to track the time-averaged Ca^2+^ levels. The addition of 100 μM Bay K8644 triggered an increase in luminescence. Then, addition of 5% Triton X-100 caused the consumption of the remaining aequorin. The upper graph shows the luminescence rate (L), the middle graph shows the % of remaining counts along the experiment (% Lmax) and the lower graph shows the Ca^2+^ levels, expressed as the ratio L/Lmax (see Materials and Methods). It is worth noting that luminescence values have no meaning in terms of Ca^2+^ levels until they are converted into ratios L/Lmax. This ratio, which is proportional to Ca^2+^ concentration [32], was independent of the total amount of functional aequorin in the sample (Suppl. Fig. 6).

**Fig. 4.**
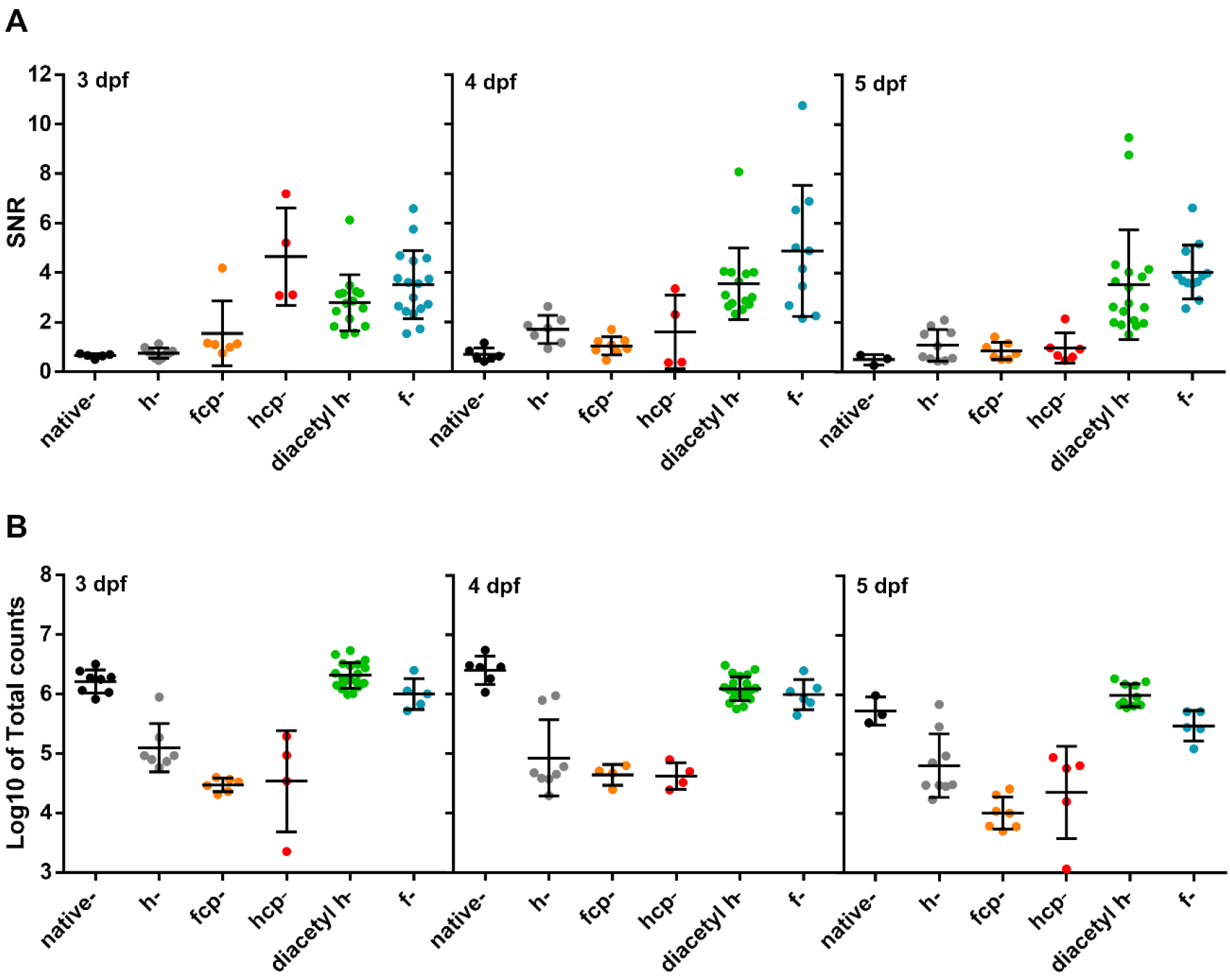
Test of CTZ analogs in 3, 4 and 5 dpf *Tg*(*cmlc2:GA*) zebrafish embryos. **A)** SNR from embryos incubated with the CTZ analogs for 2 hours. Images were acquired at 9 Hz. (*Native-CTZ* n=5, 6 and 3; *h-* CTZ n=10, 7 and 10; *fcp*-CTZ n=6, 8 and 7; *hcp-CTZ* n=4, 4 and 5; *diacetyl h*-CTZ n=15, 14 and 18; and *f*-CTZ n=17, 10 and 12, for 3, 4 and 5 dpf, respectively). Data are shown as the mean ± SD. **B)** Total counts released by the addition of 5% Triton X-100 in embryos incubated with the CTZ analogs for 2 hours. (*Native-CTZ* n=8, 6 and 3; *h*-CTZ n=7, 8 and 9; *fcp*-CTZ n=6, 4 and 7; *hcp*-CTZ n=4, 4 and 5; *diacetyl h*-CTZ n=19, 21 and 10; and *f*-CTZ n=5, 6 and 5, for 3, 4 and 5 dpf, respectively). Data are shown as the mean ± SD of log10 values.

**Fig. 5.**
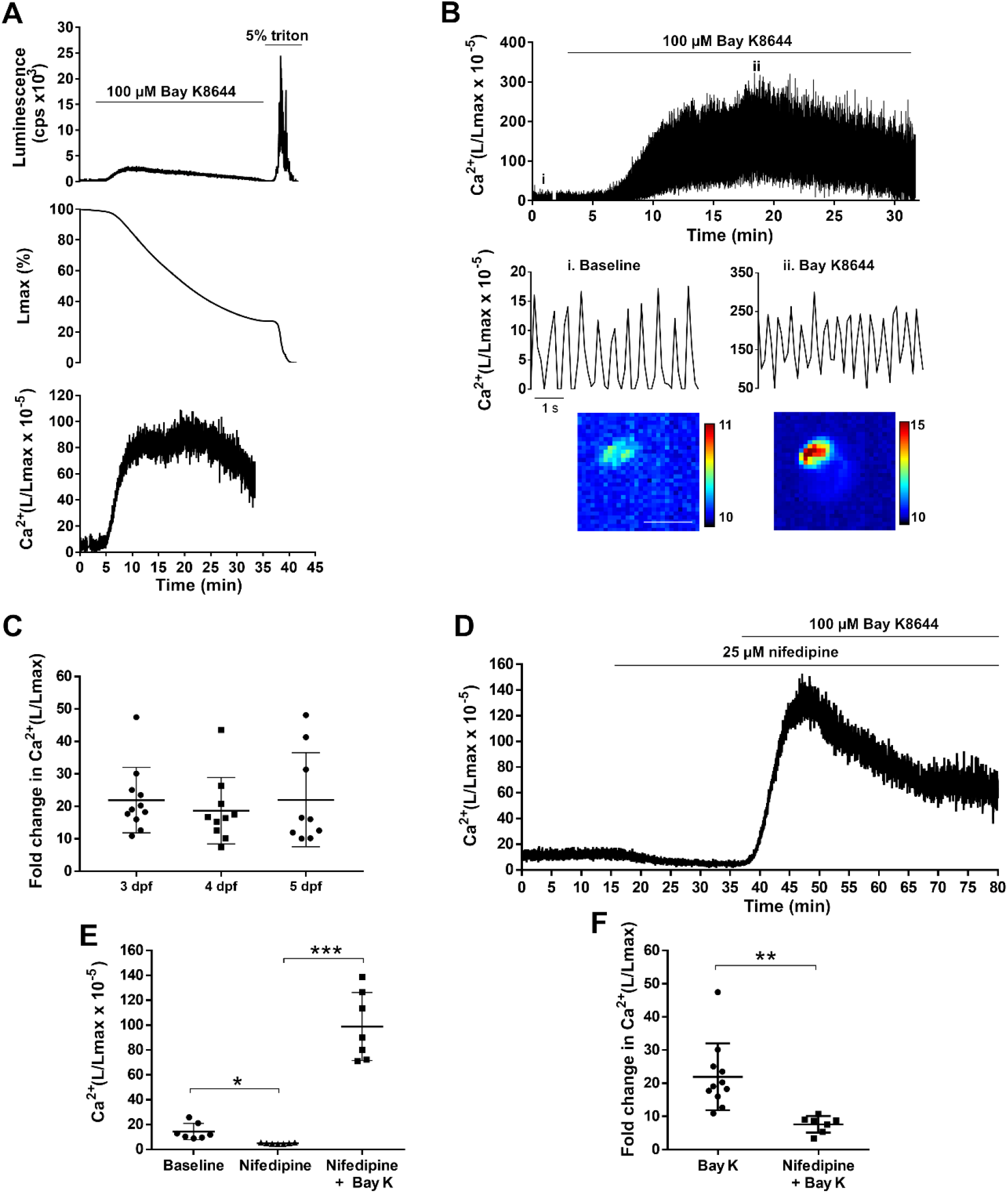
Effect of LTCC agonist and antagonist on ventricular Ca^2+^ dynamics of 3, 4 and 5 dpf *Tg*(*cmlc2:GA*) zebrafish embryos. **A)** Analysis of a bioluminescence experiment (3 dpf embryo) in which 100 μM Bay K8644 was added and Triton X-100 released all remaining counts. Upper panel: Luminescence in counts per s (L, cps); middle panel: remaining counts along the experiment (% of Lmax). Lower panel: L/Lmax, which is proportional to Ca^2+^ levels. Images were acquired at 2 Hz. **B)** Effect of 100 μM Bay K8644 on the ventricular Ca^2+^ levels. Images were acquired at 9 Hz. The lower panels zoom in the baseline (i) and 100 μM Bay K8644 (ii, min 19). The integrated luminescence images during 1 min in baseline and 100 μM Bay K8644 (min 19-20) are shown below. The scale bar represents 100 μm and the color scale indicates RLU. **C)** Fold change over basal ventricular Ca^2+^ levels (L/Lmax) of embryos treated with 100 μM Bay K8644 at 3, 4 and 5 dpf. A one-way ANOVA with Holm-Sidak *post hoc* correction for multiple comparisons and multiple *t*-tests was used. Data are shown as the mean ± SD (n=11 for 3 and 4 dpf, and n=9 for 5 dpf). **D)** Representative experiment showing the effect of 25 μM nifedipine followed by 100 μM Bay K8644 in a 3 dpf embryo. Images were acquired at 1 Hz. **E)** Effect of 25 μM nifedipine and 100 μM Bay K8644 on ventricular Ca^2+^ levels in 3 dpf embryos treated as in D). A repeated measures one-way ANOVA test was used. Data are shown as the mean ± SD (n=7). **F)** Fold change in Ca^2+^ (L/Lmax) of 100 μM Bay K 8644 in the absence (n=11) or presence (n=7) of 25 μM nifedipine. A two-tailed unpaired *t*-test was used. Data are shown as the mean ± SD. All experiments in this figure were performed with *diacetyl h*-CTZ. * p < 0.05, ** p < 0.01 and *** p < 0.001.

The effect of Bay K8644 on individual ventricular Ca^2+^ transients was studied in recordings acquired at 9 Hz (Fig. 5B). The Ca^2+^ transient amplitude gradually increased during drug addition, as well as the diastolic and systolic Ca^2+^ levels. Bay K8644 triggered a 21.9, 18.7 and 20.8-fold increase in L/Lmax at 3, 4 and 5 dpf, respectively, and no statistical difference was found between them (Fig. 5C). The solvent (0.5% DMSO) had no effect on the ventricular Ca^2+^ levels.

We investigated the effect of the LTCC antagonist nifedipine followed by the agonist Bay K8644 on ventricular Ca^2+^ levels (Fig. 5D). Nifedipine (25 μM) decreased time-averaged Ca^2+^ levels (L/Lmax 14.34 ± 6.49 before nifedipine *vs*. 4.96 ± 0.50 after nifedipine; p=0.015), an effect that was reversed by the subsequent addition of Bay K8644 (100 μM) (L/Lmax 4.96 ± 0.50 for nifedipine *vs*. 98.83 ± 27.33 after Bay K8644; p=0.0002) (Fig. 5E). The increase in L/Lmax induced by Bay K8644 was lower in the presence of nifedipine (7.64 ± 2.49 with nifedipine *vs*. 21.92 ± 10.07 without nifedipine; p=0.002) (Fig. 5F). Thus, the ability to image Ca^2+^ for extended periods allowed us to test the reversal of the effect of the LTCC antagonist by excess agonist.

### 2.5 Decreasing adrenergic tone with propranolol

The next experiment aimed to track changes in Ca^2+^ levels induced by the β-adrenergic antagonist propranolol in *Tg*(*cmlc2:GA*) embryos. Fig. 6A and B show that propranolol (100 μM) reduced Ca^2+^ levels in a 4 dpf embryo. The L/Lmax decreased 47.9 % after 30 min of exposure (55.34 ± 22.58 baseline *vs*. 27.99 ± 9.12 after propranolol; p=0.006; Fig. 6C). These results show that some adrenergic tone was present in the heart of 4 dpf embryos, influencing the time-averaged Ca^2+^ levels.

**Fig. 6.**
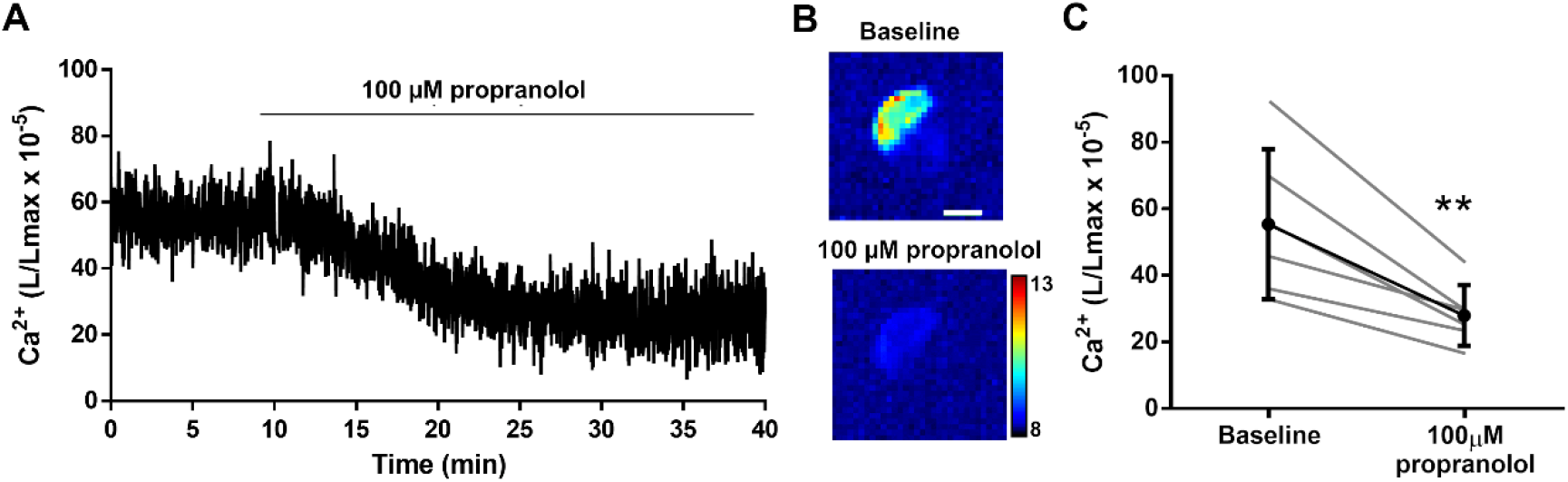
Effect of the β-adrenergic antagonist propranolol (100 μM) on the ventricular Ca^2+^ levels in 4 dpf *Tg*(*cmlc2:GA*) zebrafish embryos. **A**) Representative experiment of the effect of 100 μM propranolol on ventricular Ca^2+^ levels (L/Lmax) (1 Hz acquisition frequency). **B**) The images show the integrated luminescence from baseline (min 0-10) and propranolol (min 30-40) periods of the experiment in A). The scale bar represents 100 μm and the color scale indicates RLU. **C**) Ca^2+^ levels (L/Lmax) after 30 min of treatment with 100 μM propranolol. A two-tailed paired *t*-test was used. Data are shown as the mean ± SD (n=6). These experiments were performed using *diacetyl h*-CTZ. ** p < 0.01.

### 2.6 Ca^2+^ levels in a terfenadine-induced heart failure model

It has been reported that the treatment of 3 dpf zebrafish embryos with 10 μM terfenadine for 24 hours reproduced some features of heart failure, like heart chamber dilatation, reduced fractional shortening, arrhythmia, and apoptosis [40, 41]. Terfenadine is a potent hERG blocker and can induce prolongation of the QT interval as well as 2:1 atrioventricular block [42]. We measured Ca^2+^ levels and mechanical function in this pathological model. A decreased atrial HR was observed in the terfenadine-treated embryos with respect to control (183 ± 24 for terfenadine *vs*. 224 ± 14 bpm for the control; p<0.0001). In addition, the atrioventricular HR ratio in the terfenadine-treated embryos was 0.51 ± 0.11 (n=19), indicating a 2:1 block, which was found in 94% of the embryos. The fractional shortening was calculated with both the major and minor diameters of the ventricle (Fig. 7A). In control embryos, it was similar regardless of the diameter employed (0.36 ± 0.03 for major *vs*. 0.37 ± 0.05 for minor diameter; p=0.414). However, differences were observed in terfenadine-treated embryos (0.26 ± 0.05 for major *vs*. 0.44 ± 0.04 for minor diameter; p<0.0001) suggesting that contraction was altered (Fig. 7B) [40, 41]. We therefore examined Ca^2+^ levels by cardioluminescence. Terfenadine-treated embryos had lower time-averaged Ca^2+^ levels than the controls (Fig. 7C and D). L/Lmax values were 57 % lower in terfenadine embryos (26.01 ± 12.06 for terfenadine *vs*. 59.94 ± 13.61 for control; p=0.0003) (Fig. 7D).

**Fig. 7.**
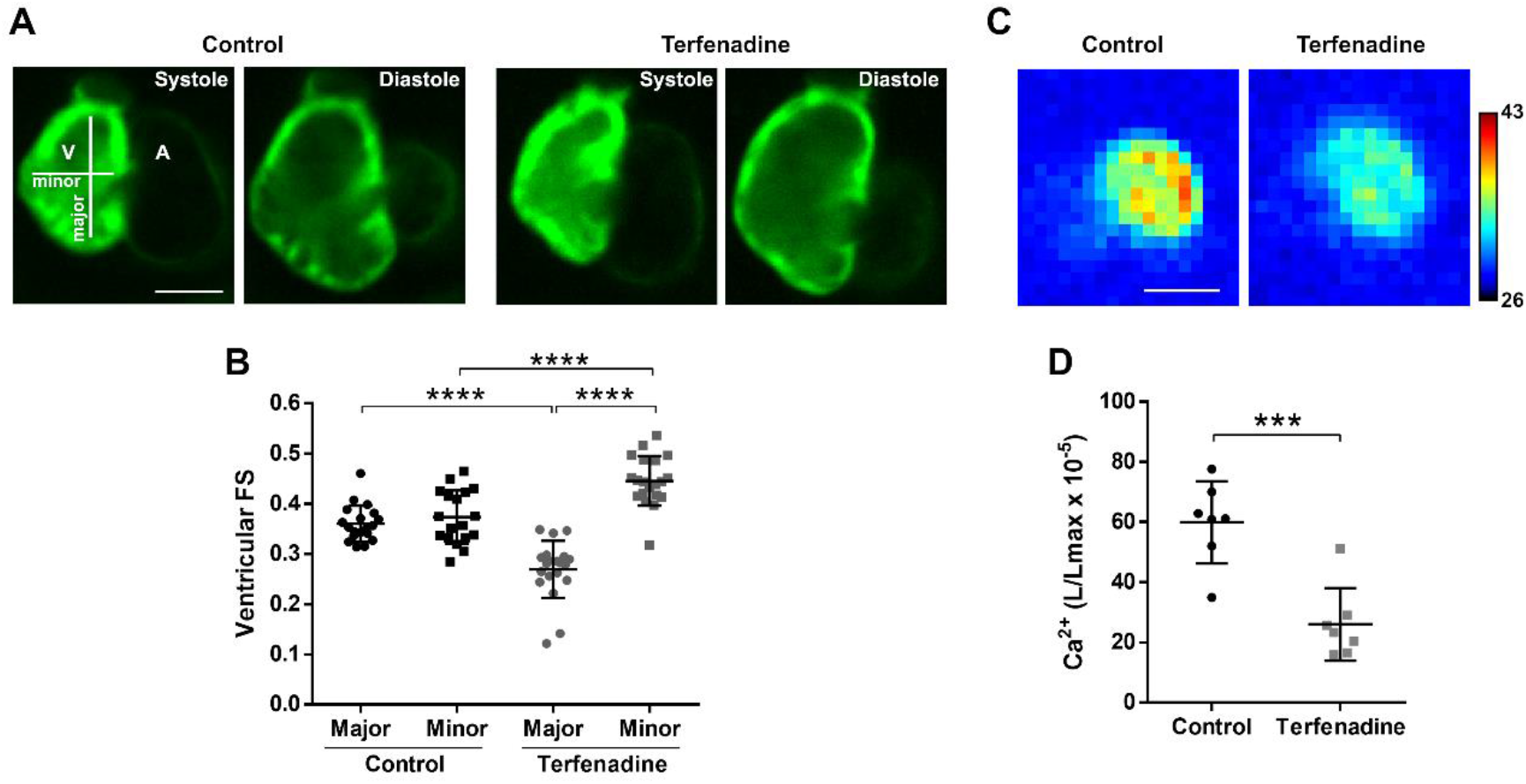
Ventricular shortening and Ca^2+^ levels in a terfenadine-induced heart failure model. *Tg*(*cmlc2:GA*) embryos (3 dpf) were treated with 10 μM terfenadine for 24 hours. **A)** The images show the ventricle in systole and diastole from representative control (0.02% DMSO) and terfenadine-treated embryos (GA fluorescence). The major and minor diameters are indicated. **B)** Ventricular fractional shortening (FS, measured with the major and minor diameters) in control and terfenadine-treated embryos. A one-way ANOVA with Holm-Sidak *post hoc* correction for multiple *t*-test comparisons was used. Data are shown as the mean ± SD (n=19 in both groups). **C)** One min integrated luminescence of control and terfenadine-treated embryos (1 Hz acquisition frequency). **D)** Ca^2+^ levels (L/Lmax) of control and terfenadine-treated embryos. A two-tailed unpaired *t*-test was used. Data are shown as the mean ± SD (n=7 in both groups). Luminescence experiments shown in C and D were performed using *diacetyl h*-CTZ. The scale bar in A and C represents 100 μm and the color scale indicates RLU. *** p < 0.001 and **** p < 0.0001.

## 3. Discussion

Here, we describe a new model for the study of Ca^2+^ physiology and pathophysiology in the embryonic zebrafish heart, the transgenic line *Tg*(*cmlc2:GA*), which reports cytosolic Ca^2+^ by bioluminescence as a complementary method to the fluorescent Ca^2+^ biosensors. Imaging live specimens by fluorescence is often limited to short duration recordings due to the phototoxicity and photobleaching that arise from strong illumination, particularly in scanning and spinning disc confocal microscopy. A notable exception is light sheet fluorescence microscopy, which has been shown to be much less harmful [20]. Fluorescence imaging of the heart usually requires stopping contraction to avoid motion artifacts [19, 21] at the cost of losing mechano-electrical feedback mechanisms. In addition, these techniques are all affected by autofluorescence of the vitello and that due to compounds like methylene blue, used in the fish water. To overcome these issues, we have used here bioluminescence imaging of aequorin fused to GFP (GA) to monitor continuously Ca^2+^ levels in experiments of up to 2 hours in 3, 4 and 5 dpf embryos, while the heart was performing its mechanical function.

Reconstitution of apoaequorin with CTZ in single-cells or *in vivo* is not generally a major problem [26, 27, 29, 30, 43]. However, in the heart newly reconstituted aequorin was rapidly consumed since Ca^2+^ is oscillating continuously. It has been previously reported that nifedipine-treated embryos completely restored the frequency and amplitude of heart Ca^2+^ transients after washout [19]. Thus, we treated embryos with nifedipine in free-Ca^2+^ medium to temporarily abolish Ca^2+^ transients during reconstitution with CTZ. No deleterious effects on heart function were observed after this procedure, which resulted in high levels of active aequorin, allowing Ca^2+^ imaging in live embryos. The duration of the bioluminescence recordings is limited by aequorin consumption. The acetyl groups in *diacetyl h*-CTZ protect it from autooxidation [39] and increased the amount of functional aequorin. Thus, the use of *diacetyl h*-CTZ enhanced the SNR and total counts, conferring long-term imaging.

We focused on the ventricular Ca^2+^ levels since atrial bioluminescence was under the limit of detection of our system. The atrium, being thinner than the ventricle, contained less amount of GA. Nevertheless, atrial bioluminescence was observed by increasing the Ca^2+^ levels with Bay K8644 or after releasing all counts with detergent. Imaging, in contrast to photometry, allowed us to quantify ventricular Ca^2+^ by setting appropriate ROIs. Since the SNR decayed at high acquisition frequencies, 9 images per s were acquired in some experiments to maintain a balance between temporal resolution and SNR. This acquisition rate sufficed to resolve individual Ca^2+^ transients in the heart, whereas lower frame rates (1-2 Hz) provided information about the steady state Ca^2+^ levels.

We verified that the *Tg*(*cmlc2:GA*) line was sensitive enough to detect increases and decreases of the Ca^2+^ levels by using an agonist (Bay K8644) and an antagonist (nifedipine) of LTCCs by modulating Ca^2+^ influx. Since images could be acquired continuously, we studied the opposite effect of these drugs on ventricular Ca^2+^ in long recordings. Ca^2+^ levels also decreased by the β-adrenergic blocker propranolol. Using fluorescent biosensors, we and others have reported a decrease in HR and Ca^2+^ levels induced by propranolol [19, 24]. The treatment of 3 dpf zebrafish embryos with the antihistamine drug terfenadine for 24 hours has been reported to induce heart failure [40, 41], involving arrhythmia and systolic disfunction, but potential changes in cytosolic Ca^2+^ were not investigated. The proarrhythmic risk of terfenadine may arise from any of its numerous effects on the cardiomyocyte electrical system: block of Na^+^ and L-type inward Ca^2+^ currents, which slows ventricular conduction and promotes non-Torsades de pointes ventricular tachycardia and fibrillation [42]; increased frequency of spontaneous Ca^2+^ release from the sarcoplasmic reticulum and enhanced NCX spontaneous currents [44]; and block of hERG with QT prolonging effects [42]. In zebrafish, the prolongation of the repolarization has been shown to result in atrioventricular block [45]. Our results showed a significant decrease in the ventricular Ca^2+^ levels in terfenadine-treated embryos that can be associated with a decrease in the HR induced by the 2:1 atrioventricular block. We reported that the Ca^2+^ levels decreased concomitantly with a decrease of the HR in zebrafish embryos treated with propranolol [24]; in addition, Zhang et al. reported a decrease in Ca^2+^ when zebrafish ventricular myocytes were paced at a slower rate [6].

Since the bioluminescence reaction of aequorin is triggered by three Ca^2+^ ions [46], the steepness of the Ca^2+^-response curve of aequorin confers an excellent SNR. However, in Ca^2+^ microdomains near sarcolemmal Ca^2+^ channels or ryanodine receptors on the sarcoplasmic reticulum surface (Ca^2+^ sparks) [47], a small fraction of total aequorin may be exposed to very high Ca^2+^ concentration. Thus, L/Lmax signal may be dominated by these microdomains and not represent the averaged cytoplasmic Ca^2+^ levels. In fact, the experiments with Bay K8644, which induced a 20-fold change in L/Lmax, suggest the existence of such microdomains.

Luminescence may be close to noise levels when cardiac Ca^2+^ levels decrease either by drugs or in a pathological model. Increasing the Ca^2+^ affinity of the photoprotein by mutations in apoaequorin or with appropriate CTZ analogs can overcome this limitation [31] at the cost of speeding up the aequorin consumption, as was observed with the high Ca^2+^ affinity *hcp-CTZ* (Fig. 4B). Alternatively, lowering the imaging acquisition rate increases the SNR, as was shown with both nifedipine and propranolol, but individual Ca^2+^ transients may not be resolved. This highlights the versatility of GA bioluminescence to monitor oscillatory Ca^2+^ transients or time-averaged levels.

In conclusion, we have constructed a transgenic zebrafish line, *Tg*(*cmlc2:GA*), expressing the bioluminescent Ca^2+^ biosensor GA in the heart, and devised a protocol for aequorin reconstitution. Ventricular Ca^2+^ dynamics were imaged continuously in beating hearts for up to two hours in 3-5 dpf embryos. The time-averaged Ca^2+^ levels and individual Ca^2+^ transients were monitored and, as a proof-of-concept, we studied changes induced by drugs acting on LTCC and sympathetic input. *Tg*(*cmlc2:GA*) embryos also revealed a decrease in Ca^2+^ levels in a model of heart failure induced by terfenadine. Fluorescence and luminescence imaging are orthogonal techniques to interrogate pathophysiological processes, but cardioluminescence avoids light toxicity and allows monitoring Ca^2+^ for longer periods of time.

## 4. Materials and Methods

### 4.1 Zebrafish husbandry

Fish used in this study were housed under standard conditions as previously described [29]. All animal procedures were carried out in accordance with institutional and national ethical and animal welfare guidelines (approval identification code 900823 dated 30 May 2016, Consejería de Agricultura, Medio Ambiente y Desarrollo Rural, JCCM, Spain; and Max Planck Gesellschaft and the ethics committee for Regierungspräsidium Darmstadt, Germany).

### 4.2 Generation of Tg(cmlc2:GA) zebrafish line

The Ca^2+^ biosensor GA [33] was cloned into the *pT2A-Tol2-cmlc2* transposon vector using *XhoI* and *Eco*RI restriction sites. To generate stable transgenic zebrafish, *tol2-cmlc2:GA* plasmid (12.5 ng/μL) was co-injected with transposase mRNA (12.5 ng/μL) in Tübingen zebrafish fertilized eggs. Injected embryos (F0) were screened by fluorescence for GA expression in the heart and grown to adulthood. The adult F_0_ generation was outcrossed to wild-type zebrafish to identify founders with insertions in the germline. F2 *Tg*(*cmlc2:GA*) heterozygous embryos were used throughout the study.

### 4.3 Synthesis of diacetyl CTZ-h

CTZ-*h* was synthesized using the procedures in the previous report [48]. *h-*CTZ (73.2 mg, 0.180 mmol) and DMAP (60.0 mg, 0.491 mmol) in acetic anhydride (2.50 mL, 26.4 mmol) were stirred overnight at 20 °C under a N2 atmosphere. After removal of all the volatiles, the residue was dissolved in ethyl acetate and washed with 2 M HCl, saturated aqueous NaHCO_3_, and brine, with the organic layer then dried over Na2SO4, filtered, and evaporated in vacuo. The residue was purified by silica gel column chromatography using 5-10% ethyl acetate in dichloromethane as eluent to afford *diacetyl h*-CTZ as a yellow-brown solid (64.3 mg, 0.131 mmol, 73%). 1H NMR (500 MHz, CDCl_3_): δ 7.87 (d, *J* = 9.0 Hz, 2H), 7.73 (s, 1H), 7.84 (d, *J* = 7.5 Hz, 2H), 7.30-7.20 (m, 8H), 7.15 (d, *J* = 9.0 Hz, 2H), 4.60 (s, 2H) 4.18 (s, 2H) 2.29 (s, 3H), 2.12 (s, 3H). MS (ESI+) Calcd for [M+H]^+^, 492.19; found 492.19.

### 4.4 Aequorin reconstitution with coelenterazine

A stock of *diacetyl h*-CTZ was prepared at 7.4 mM in dimethyl sulfoxide (DMSO). The stocks of *native*-, *hcp*-, *fcp*-, *h*- and *f*-CTZ analogs (Biotium, Fremont, CA, USA) were prepared at 5 mM in methanol and 5 μL aliquots were stored at −80°C. All CTZs were used at 50 μM final concentration. For aequorin reconstitution, *Tg*(*cmlc2:GA*) zebrafish embryos at 3, 4 and 5 dpf were rinsed in zero Ca^2+^ E3 medium (5 mM NaCl, 0.17 mM KCl, 0.33 mM MgSO4, 0.002% methylene blue, pH 7.4 in double distilled H_2_O). Subsequently, embryos were incubated in 1 mL of zero Ca^2+^ E3 medium containing 25 μM nifedipine for 30 min at room temperature in the dark (*nifedipine period* in Fig. 2B). Embryos whose heart was completely stopped after this treatment were washed out 5 times and incubated in zero Ca^2+^ E3 medium containing 50 μM CTZ for 2 hours at room temperature in the dark (*CTZ incubation period* in Fig. 2B). Finally, the embryos were incubated in complete E3 medium (5 mM NaCl, 0.17 mM KCl, 0.33 mM MgSO_4_, 0.33 mM CaCl_2_, 0.002% methylene blue, pH 7.4 in double distilled H_2_O) for 30 min at 28.5°C (*heart recovery period* in Fig. 2B).

### 4.5 Bioluminescence imaging

After aequorin reconstitution, *Tg*(*cmlc2:GA*) embryos were embedded in 100 μL of 0.3% low melting agarose prepared in E3 medium and transferred to an 8-well glass bottom plate (ibidi, Gräfelfing, Germany). When the agarose solidified, 100 μL of E3 medium was added and a small portion of agarose surrounding the heart of the embryos was cut out to improve diffusion of drugs and Triton X-100. For the signal-to-noise ratio (SNR) calculation, bioluminescence images were acquired at 9 Hz for 1 min. For longer recordings, drugs were added after 1-10 min basal tracking. At the end of every recording Triton X-100 (5%) was added to release all luminescence counts from the remaining active aequorin. Bioluminescence images were acquired as previously described [1] with a custom-built microscope housed in a light-tight box to maintain complete darkness during imaging. Images were acquired continuously in 16 bits with 4 x 4 binning, 255 EM gain, at a rate of 25, 17, 12, 9, 2 or 1 Hz (frames/s) with an EM-CCD camera (Hamamatsu Photonics, Hamamatsu, Japan). The spatial resolution of the images was 12.8 μm x 12.8 μm/pixel and the total field of view was 1638 μm x 1638 μm.

### 4.6 GCaMP fluorescence imaging

GCaMP fluorescence imaging was carried out at the Max Planck Institute for Heart and Lung Research (Bad Nauheim, Germany). *Tg*(*cmlc2:GCaMP*)^*s878*^ adult zebrafish were outcrossed to wild-type strain and fertilized eggs at 1-cell stage were injected with 2 ng of the morpholino oligomer *tnnt2a* (5’-CATGTTTGCTCTGATCTGACACGCA-3’). Embryos at 24 hours post-fertilization were placed in 0.003% N-phenylthiourea to prevent pigmentation. For nifedipine titration, 3 dpf embryos were incubated in E3 medium or zero Ca^2+^ E3 medium containing 10, 25 or 100 μM nifedipine for 30 min. Then, embryos were embedded in 1% low melting point agarose and transferred to an 8-well glass bottom plate (ibidi). To study the recovery of Ca^2+^ dynamics with GCaMP, the aequorin reconstitution protocol described above without CTZ was applied to 3 dpf embryos. Fluorescent images were acquired at a rate of 200 Hz with a CSU X1 spinning disc confocal microscope (Carl Zeiss, Oberkochen, Germany) equipped with a Hamamatsu ORCA Flash4.0 sCMOS camera (Hamamatsu Photonics, Japan) in 16 bits with 2 x 2 binning.

### 4.7 Pharmacological treatments

Chemical compounds were dissolved in DMSO to prepare stocks of 10 mM nifedipine (Sigma-Aldrich N7634, Darmstadt, Germany), 20 mM Bay K8644 (Tocris 1544), 10 mM propranolol (Sigma-Aldrich P0884, Darmstadt, Germany), 50 mM terfenadine (Tocris 3948) and 7.5% N-phenylthiourea (Sigma-Aldrich, Darmstadt, Germany).

### 4.8 Heart failure induced by treatment with terfenadine

Embryos at 24 hours post-fertilization were placed in 0.003% N-phenylthiourea in E3 medium to prevent pigmentation. At 3 dpf, embryos were transferred to a 6-well plate, 10 embryos per well, with 5 mL of N-phenylthiourea solution containing 10 μM terfenadine or 0.02% DMSO, for 24 hours. For bioluminescence experiments, the aequorin reconstitution was done during the last 3 hours of treatment, and terfenadine or DMSO were maintained throughout.

### 4.9 Data analysis

Videos in TIFF format were analyzed in Fiji-imageJ (U.S. National Institutes of Health, Bethesda, Maryland, USA) [49]. For GCaMP image analysis, regions of interest (ROI) were drawn in the atrium and in the ventricle to obtain mean intensity values. An exponentially weighted moving average smoothing with a smoothing factor of 0.7 was applied and data was transformed into ΔF/F_0_= (F_t_ – F_0_)/F_0_; where F_t_ is the fluorescence at a given time and F_0_ is the minimum diastolic fluorescence value. For characterization of the Ca^2+^ transients, ΔF/F0 data were analyzed with Clampfit 10.7 (Molecular Devices, San José, CA, USA) to determine peak amplitude ((Fsystole-F_0_)/F_0_), rise time 10% to 90% and decay time 90% to 10%. For bioluminescence image analysis and calculation of SNR, a time-projection of the stack was performed to draw the ROIs over the ventricle and atrium (*Signal*). Then, ROIs of identical size were placed in 6 different locations far from the embryo to obtain the average background intensity values and their SD. The SNR for each frame was calculated as follows: SNR = (Signal – Background) / SD Background.

Luminescence signal (in relative light units, RLU) was transformed into Luminescence rate (L, in counts per second). The total counts were obtained as the sum of luminescence (RLU) along the experiment. Finally, Lmax was calculated as the remaining counts at each time of the experiment. Ventricular fractional shortening was calculated from end-diastolic (D_diastole_) and end-systolic (D_systole_) diameters of the ventricular cavity: fractional shortening = (D_systole_-D_diastole_)/D_diastole_ in transmitted light or fluorescence images, as indicated. Images were acquired at a rate of 50 Hz with a wide-field fluorescence microscope (DMIRE-2, Leica Microsystems, Wetzlar, Germany) as previously described [24].

### 4.10 Statistics

The Shapiro-Wilk statistic was used to test for normality. Differences between two groups were analyzed using the unpaired or paired two-tailed *t*-test, as indicated. One-way ANOVA and two-way ANOVA with Holm-Sidak *post hoc* correction for multiple comparisons and multiple *t*-tests were used when indicated. Correlation between two datasets was analyzed using linear regression. Datasets of total counts were transformed into log10 values. Data are presented as mean ± SD and p < 0.05 was considered statistically significant. * p < 0.05, ** p < 0.01, *** p < 0.001, **** p < 0.0001. N indicates the number of embryos used per dataset. Data were analyzed in Graphpad Prism 6 (GraphPad Software, Inc.; La Jolla, CA, USA).

## Acknowledgments

This research was funded by the Ministry of Science, Innovation and Universities, Spain (BFU2015-69874-R and PID2019-111456RB-100, co-funded by EU FEDER-ERDF), by Consejería de Educación, Cultura y Deportes, Junta de Comunidades de Castilla-La Mancha (SBPLY/19/180501/000223, co-funded by EU FEDER-ERDF) and grants for research groups from University of Castilla-La Mancha (UCLM) (2019-GRIN-27019 and 2020-GRIN-29186, co-funded by EU FEDER-ERDF). J.S.-A. held a predoctoral fellowship from UCLM. M.V. obtained a fellowship from University of Castilla-La Mancha to visit the Max Planck Institute for Heart and Lung Research, Bad Nauheim, Germany.

We thank Didier Stainier for use of the *tg*(*cmlc2:GCaMP*)^*s878*^ line at the Max Planck Institute for Heart and Lung Research. We thank Pierre Vincent and Eduardo Nava for valuable suggestions, and Carmen Cifuentes for expert technical assistance.

## Conflicts of Interest

The authors declare no conflict of interest. The funders had no role in the design of the study; in the collection, analyses, or interpretation of data; in the writing of the manuscript, or in the decision to publish the results.

## Abbreviations

CTZ: coelenterazine
DMSO: dimethyl sulfoxide
dpf: days post-fertilization
FS: fractional shortening
GA: GFP-Aequorin
GFP: green fluorescent protein
HR: heart rate
LTCC: L-type calcium channel
RLU: relative luminescence unit
ROI: region of interest
SNR: signal-to-noise ratio.

## Author Contributions

M.V. generated the transgenic zebrafish line. M.M. and K.K. synthesized diacetyl coelenterazine-*h*. B.D. and J.L. set up the bioluminescence microscope. M.V., B.D. and J.L. designed the experiments. M.V. performed the bioluminescence experiments. J.S. and A.M. performed the fluorescence experiments with GA. M.V. and M.M. C. performed the GCaMP experiments. M.V., B.D. and J.L. analyzed the data and wrote the manuscript. B.D. and J.L. obtained funding.

## Notes

### Competing Interest Statement

The authors have declared no competing interest.

