## Supplementary material for "Cardioluminescence in transgenic zebrafish embryos: a Ca^2+^ imaging tool to study drug effects and pathological modeling"

### Supplementary Figures:

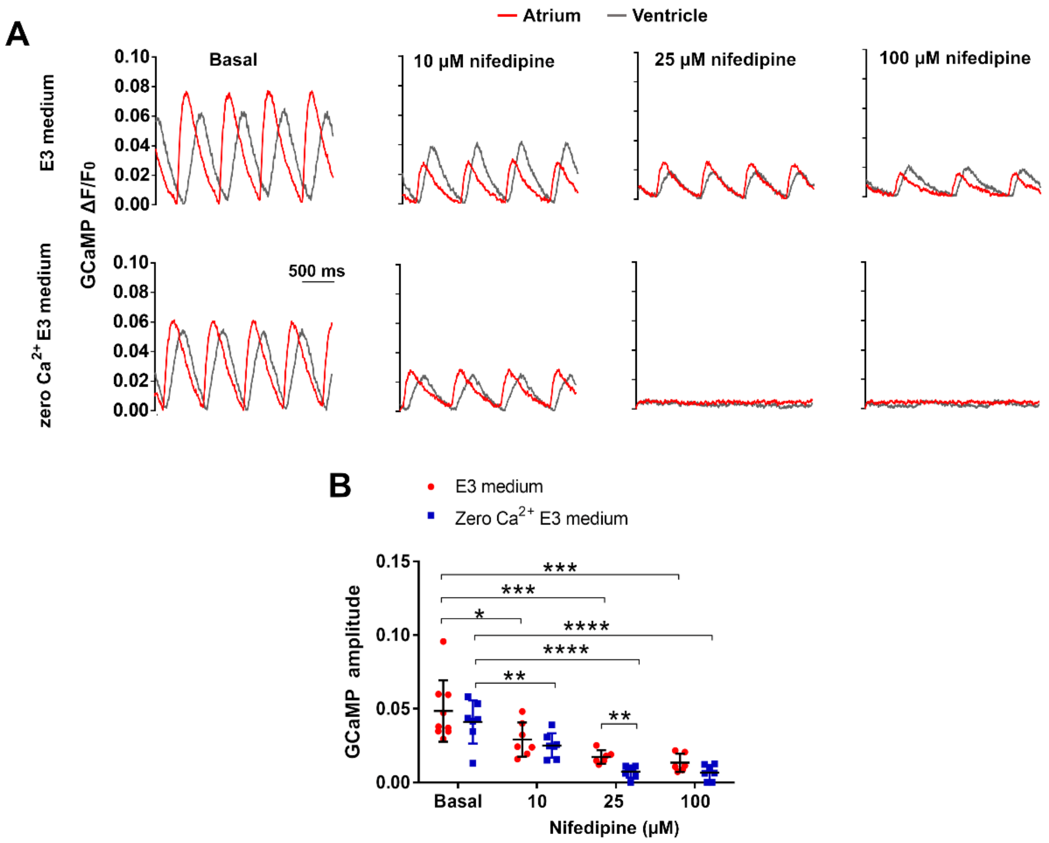


**Suppl. Fig. 1. Dose-response of nifedipine on $Ca^{2+}$ transients measured with GCaMP.** Three concentrations of nifedipine were tested in the absence or presence of $Ca^{2+}$ in E3 medium in 3 dpf *Tg(cmlc2:GCaMP)*$^{s878}$ embryos. **A)** Representative $Ca^{2+}$ transients with 10, 25 and 100 μM nifedipine for 30 min in complete E3 medium (upper panel) or in zero $Ca^{2+}$ E3 medium (lower panel). The fluorescence images were acquired at 200 Hz. **B)** Dose-response of nifedipine on ventricular $Ca^{2+}$ transient amplitude in both media. A two-way ANOVA with Holm-Sidak *post hoc* correction for multiple comparisons and multiple *t*-tests was used. Data are shown as the mean ± SD. E3 medium (basal n=9, 10 μM n=7, 25 μM n=6, 100 μM n=6) and zero $Ca^{2+}$ E3 medium (basal n=7, 10 μM n=7, 25 μM n=7, 100 μM n=7). * $p < 0.05$, ** $p < 0.01$, *** $p < 0.001$, **** $p < 0.0001$.

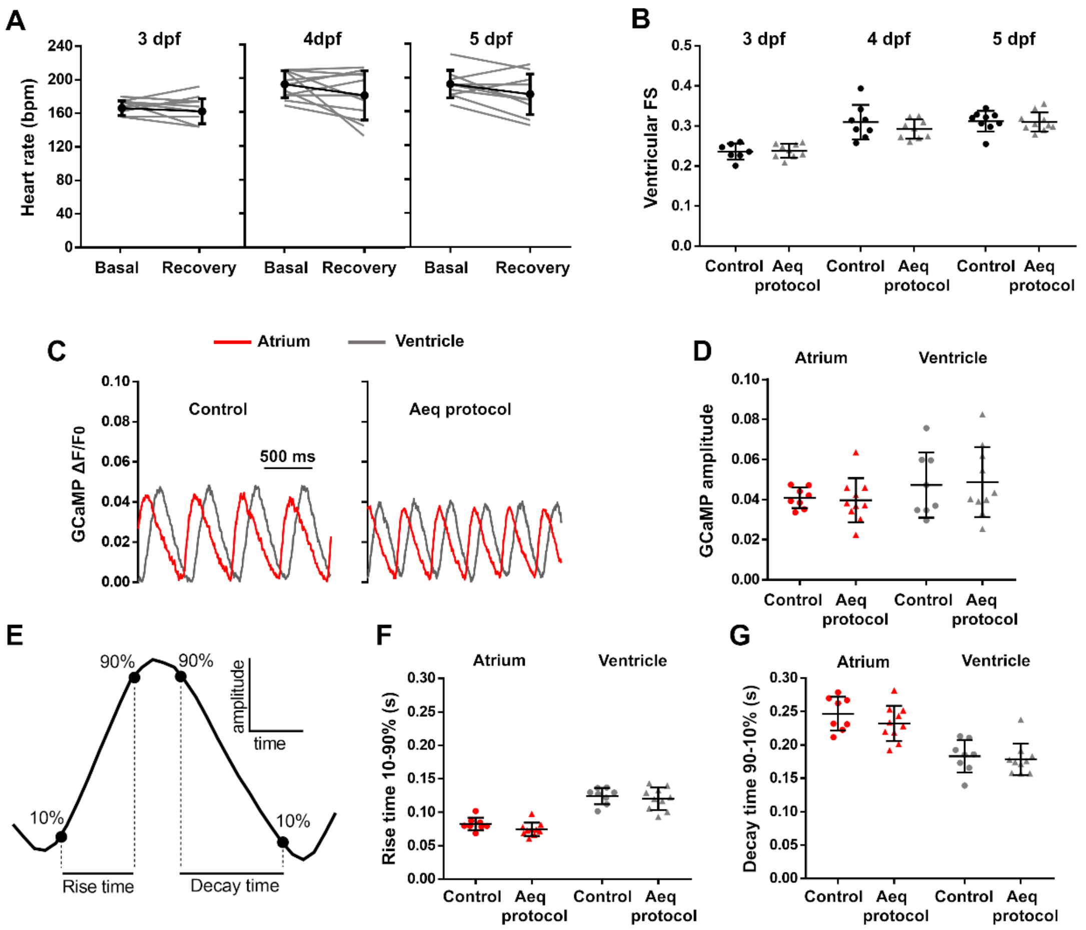


**Suppl. Fig. 2. Restoration of the heart mechanical function and the $Ca^{2+}$ transients after the recovery period of the aequorin reconstitution protocol. A)** HR during the basal and recovery periods in 3, 4 and 5 dpf *Tg*(*cmlc2:GA*) zebrafish embryos. A two-tailed paired *t*-test was used. Data are shown as the mean ± SD (n=11 for 3 and 4 dpf; n=10 for 5 dpf). The temperature was held at 27.5ºC. **B)** Ventricular fractional shortening (FS) from untreated embryos (control group) and the aequorin reconstitution group in 3, 4 and 5 dpf *Tg*(*cmlc2:GA*) embryos. A two-tailed unpaired *t*-test was used. Data are shown as the mean ± SD (control n=7 for 3 dpf, n=8 for 4 dpf and n=9 for 5 dpf; Aeq protocol n=9 for 3 and 4 dpf and n=10 for 5 dpf). **C)** $Ca^{2+}$ transients from representative control and Aeq protocol 3 dpf *Tg*(*cmlc2:GCaMP*)$^{s878}$ embryos. **D)** $Ca^{2+}$ transient amplitude in atrium and ventricle in untreated embryos (control) and Aeq protocol groups. **E)** Schematic representation of the $Ca^{2+}$ transient showing the 10-90% rise time and 90-10% decay time. **F)** Rise time and **G)** decay time in atrium and ventricle from untreated embryos (control) and Aeq protocol groups. A two-tailed unpaired *t*-test was used. Data are shown as the mean ± SD (n=8 for control and n=10 for Aeq protocol groups for D, F and G). No statistical differences were found in any of the groups ($p > 0.05$).

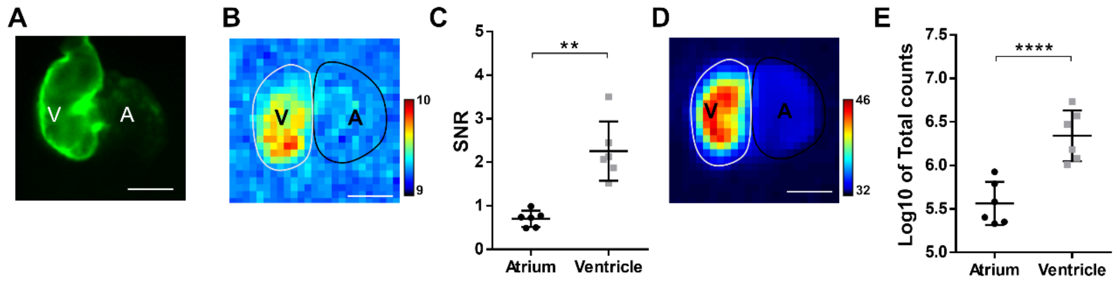


**Suppl. Fig. 3. Luminescence in the atrium and the ventricle of 3 dpf *Tg(cmlc2:GA)* embryos subjected to the aequorin reconstitution protocol. A)** Image of GFP fluorescence in the heart showing the atrium (A) and ventricle (V). **B)** Image of integrated luminescence for 1 min. Images were acquired at 9 Hz. **C)** SNR of the atrium and the ventricle. A two-tailed paired *t*-test was used. Data are shown as the mean ± SD (n=6). **D)** Image of the luminescence integrated during the whole experiment. Images were acquired at 9 Hz. **E)** Total counts released in the atrium and in the ventricle. A two-tailed paired *t*-test was used. Data are shown as the mean ± SD (n=6). *Diacetyl h*-CTZ was used for reconstitution. Scale bar in A, B and D represents 100 µm and the color scale indicates RLU. ** p < 0.01, **** p < 0.0001.

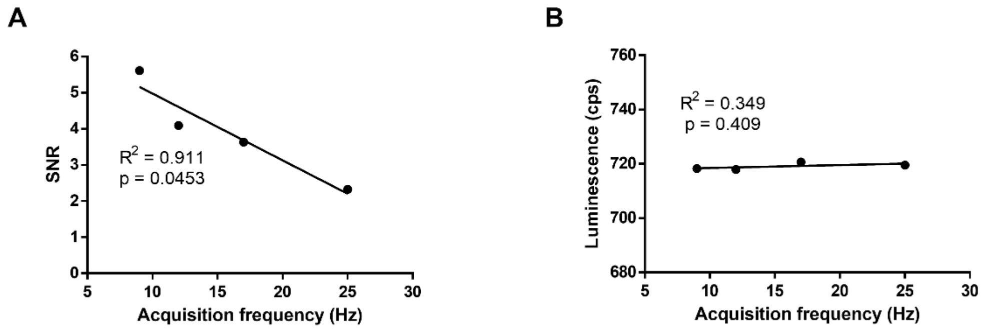


**Suppl. Fig. 4. Dependency of SNR and Luminescence rate on the image acquisition frequency.** Correlation between the image acquisition frequency (9, 12, 17 and 25 Hz) and the SNR (**A**) and the luminescence emission rate (**B**) in a 3 dpf *Tg(cmlc2:GA)* zebrafish embryo reconstituted with *diacetyl h*-CTZ.

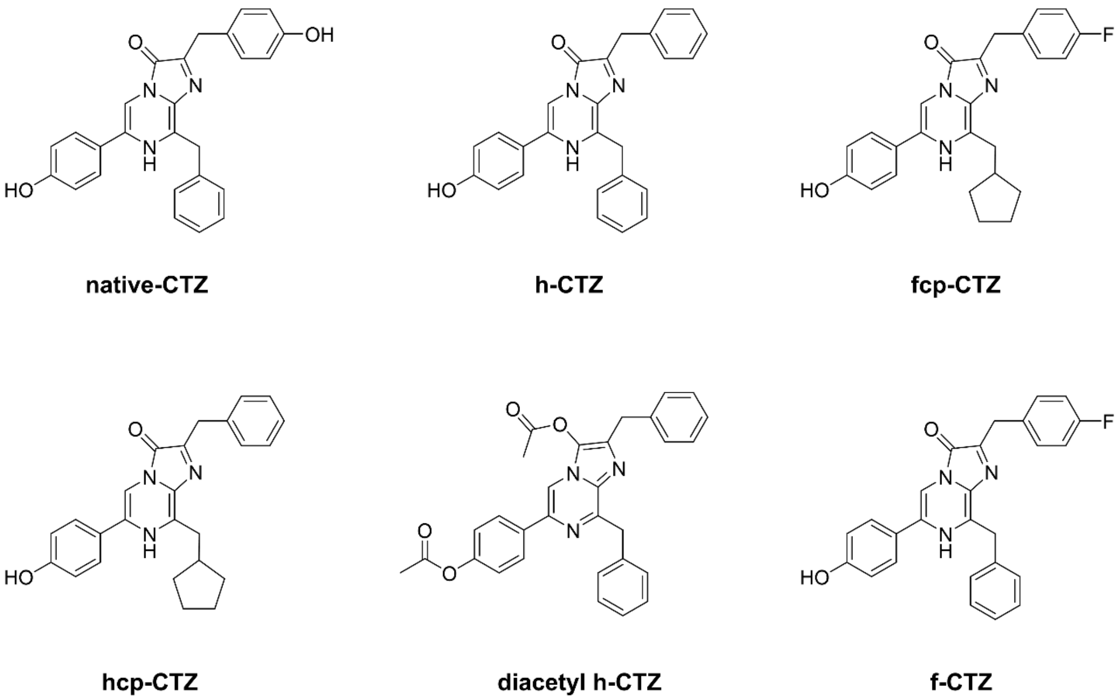


**Suppl. Fig. 5. Structure of the coelenterazines used.**

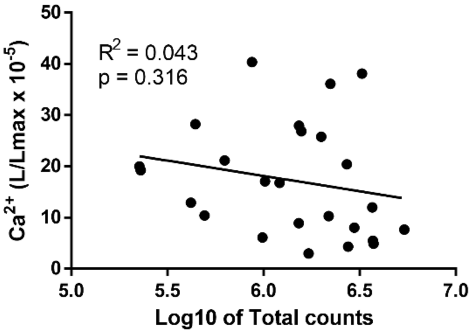


**Suppl. Fig. 6. Correlation of the basal L/Lmax values versus the total counts in 3 dpf *Tg(cmlc2:GA)* zebrafish embryos**. These experiments were performed using *diacetyl h*-CTZ. A linear regression test was used (n=25).
